# The neural processes underpinning flexible semantic retrieval in visual and auditory modalities

**DOI:** 10.1101/2025.06.22.660970

**Authors:** Ximing Shao, Meichao Zhang, Xiuyi Wang, Andre Gouws, Rebecca L. Jackson, Jonathan Smallwood, Katya Krieger-Redwood, Elizabeth Jefferies

## Abstract

Contemporary accounts of semantic cognition propose that conceptual knowledge is supported by a heteromodal conceptual store and controlled retrieval processes. However, it remains unclear how the neural basis of semantic control varies across modalities. Recent models of cortical organization suggest that control networks are distributed along a unimodal-to-heteromodal cortical gradient, with the semantic control network (SCN) located in more heteromodal cortex than the domain-general multiple demand network (MDN). We used fMRI to examine how these networks respond to semantic control demands in visual and auditory tasks. Participants judged the semantic relatedness of spoken and written word pairs. On half of the trials, a task cue specified the semantic feature to guide retrieval; on the remaining trials, no such cue was given. The SCN showed greater activation when task knowledge was available, consistent with a role in the top-down control of semantic retrieval across modalities. In contrast, the MDN showed greater activation for spoken words, likely reflecting increased demands in speech perception. These findings demonstrate a dissociation between control networks, with SCN involvement modulated by task structure and MDN activity influenced by input modality.

## 1. Introduction

Semantic cognition recruits a heteromodal network that shows common responses across spoken and written words, as well as non-verbal meaningful materials such as pictures (Jefferies, 2013; Lambon Ralph, Jefferies, Patterson, & Rogers, 2017). The Controlled Semantic Cognition account proposes that distinct brain areas within this network have different functions (Jefferies, 2013; Lambon Ralph, Jefferies, Patterson, & Rogers, 2017; Patterson, Nestor, & Rogers, 2007; Whitney et al., 2011): bilateral anterior temporal lobe (ATL) acts as a hub supporting long-term conceptual representation (Patterson, Nestor, & Rogers, 2007), while a left-lateralised control network interacts with this hub to produce flexible conceptual retrieval that can be focussed on unusual aspects of knowledge or goal-relevant features when required (Whitney et al., 2011; Noonan et al., 2013; Jackson 2021). Both the representation system that supports long-term concepts and the control system appear to be heteromodal. Patients with semantic dementia have consistent semantic deficits across different input and output systems and different types of tasks, suggesting that they have degradation of ‘central’ multimodal concepts (Patterson, Nestor, & Rogers, 2007). In contrast, people with semantic aphasia have deregulated semantic cognition affecting both verbal and non-verbal semantic tasks, such as object use (Corbett, Jefferies, & Lambon Ralph, 2011); they have difficulty controlling retrieval such that it is appropriate for the task or context, especially when the knowledge that is required is weakly encoded or subject to competition from irrelevant meanings (Jefferies and Lambon Ralph, 2006; Jefferies, 2013). Neuroimaging studies also show common recruitment of semantic regions across tasks involving spoken and written words and pictures (Binder et al., 2011; Humphreys et al., 2015; Patterson, Nestor, & Rogers, 2007).

While this research points to the existence of heteromodal semantic regions, it does not preclude the possibility that there are differences between modalities within this network. Contemporary views of cortical organisation (Huntenburg, Bazin, & Margulies, 2018; Lambon Ralph, Jefferies, Patterson, & Rogers, 2017; Margulies et al., 2016; Smallwood et al., 2021) suggest that heteromodal areas, including default mode and fronto-parietal control regions, are located relatively far from unimodal regions along the cortical surface. This spatial separation of heteromodal and unimodal cortex is thought to be a key dimension of functional cortical organisation and is captured by the ‘principal gradient’, which explains the most variance in whole-brain patterns of intrinsic connectivity. Even heteromodal semantic areas such as the anterior temporal lobes can show graded functional variation driven by the relative proximity of different subregions to converging sensory motor inputs (Lambon Ralph, Jefferies, Patterson, & Rogers, 2017). Heteromodal regions that are closer to specific primary systems might show stronger effects of modality for similar reasons (cf. Wang et al., 2023).

In addition, recent research suggests that multiple, partially dissociable control networks contribute to semantic cognition (Chiou et al., 2023; Davey et al., 2016; Wang et al., 2020; Gao et al., 2021); these control networks might show differences across modalities. The multiple-demand network (MDN), centred on inferior frontal sulcus and intraparietal sulcus, is thought to support goal maintenance and selection of relevant representations across domains (Duncan 2010; Duncan et al., 2020; Fedorenko, Duncan, & Kanwisher, 2013). In contrast, the semantic control network (SCN), with peaks in left inferior prefrontal and posterior temporal cortex, responds more selectively to control-demands within semantic tasks (Badre & Wagner, 2007; Chiou, Humphreys, Jung, & Lambon Ralph, 2018; Davey et al., 2016; Jackson 2021; Jefferies, 2013; Lambon Ralph et al., 2017; Noonan et al., 2010; Noonan et al., 2013). These networks have distinct topographies: semantic control processes are highly left-lateralised (Davey et al., 2016; Gonzalez-Alam et al., 2019; Whitney et al., 2011), while MDN is a bilateral network (Duncan 2010; Duncan et al., 2020; Fedorenko, Duncan, & Kanwisher, 2013). Moreover, SCN is sandwiched between DMN and multiple demand network (MDN) on the cortical surface, and SCN is closer to the heteromodal end of the principal gradient compared to MDN (Chiou et al., 2023; Davey et al., 2016; Wang et al., 2020), suggesting that MDN might show stronger effects of input modality. Within left inferior frontal gyrus (LIFG) and dorsomedial prefrontal cortex (DMPFC), there may be functional gradients (Diveica et al., 2023; Jung, Ralph, & Jackson, 2022; Wang et al., 2020), with more anterior and ventral aspects showing stronger connectivity with default mode network and more selective recruitment in semantic tasks, while more posterior regions support demanding sensory-motor as well as cognitive tasks – suggesting they might show greater effects of input modality (Krieger-Redwood et al., 2015; Jackson 2021).

In this study, we examined how semantic control processes interact with input modality by manipulating task knowledge during spoken and written word judgements. Prior work has shown that top-down control over semantic retrieval recruits the semantic control network (SCN), particularly left inferior frontal gyrus (LIFG), in the visual domain (Zhang et al., 2021). We extended this approach to the auditory modality to test whether the SCN supports controlled retrieval in a modality-independent fashion, and to explore how it differs from the domain-general multiple demand network (MDN) in response to semantic task demands. Participants judged whether word pairs presented either visually or aurally were semantically related via thematic (e.g., *dog–collar*) or taxonomic (e.g., *dog–chipmunk*) links. On half the trials, a cue indicated the relevant relation type in advance, enabling goal-directed semantic retrieval; on the remaining trials, no such information was provided. This design allowed us to examine how task knowledge and input modality influence engagement of distinct control networks. A gradient-based view of cortical organisation suggests that the SCN, situated in heteromodal cortex, should support controlled semantic retrieval across modalities when goals are specified in advance. In contrast, the MDN, located closer to unimodal sensorimotor cortex, may be more sensitive to modality-specific processing demands. Recognising isolated spoken words is inherently more ambiguous than reading written text, particularly in the acoustically challenging MRI environment. We therefore expect MDN activity to reflect perceptual effort more than semantic control demands. By orthogonally manipulating task structure and modality, this study tests how domain-specific and domain-general control processes are embedded within the brain’s large-scale functional architecture.

## 2. Methods

### 2.1. Design

Two groups of participants performed semantic judgements to visual or spoken words, with and without the opportunity for top-down control over retrieval. Both groups were presented with two words in succession; we manipulated top-down control by informing participants in advance about the kinds of semantic features that would be relevant to the semantic decision on half of the trials. The data from participants tested on written words were previously published (Zhang et al., 2021) and we collected new data using the same stimuli and paradigm, and employing the same scanner and imaging sequence, to examine similarities and differences between these two input modalities (slight adaptations to the methods for auditory inputs are outlined below).

### 2.2. Participants

For the auditory experiment, 32 students were recruited (mean age = 21.9 ± 3.91 years, 5 males). Five participants were excluded from data analysis due to low accuracy on the semantic decision task (*Mean ± SD* = 57.5% ± 5.69%). Consequently, 27 participants were included in the final analysis. For the visual experiment (Zhang et al., 2021), 32 students were recruited (mean age = 20.6 ± 1.52 years, 5 males). One participant was excluded from data analysis due to low accuracy on the semantic decision task and therefore 31 participants were included in the final analysis. All participants across both groups were right-handed native English speakers, and had normal or corrected-to-normal vision. None of them had any history of neurological impairment, diagnosis of learning difficulty or psychiatric illness.

A separate sample of 176 participants (mean age = 20.57, 114 females) who completed resting-state fMRI, was used to examine the intrinsic connectivity of regions identified in task contrasts. For all samples, ethical approval was obtained from the Research Ethics Committees of the Department of Psychology and York Neuroimaging Centre, University of York. All participants provided written informed consent prior to taking part and received monetary compensation or course credit for their time.

### 2.3. Materials

In both the visual and auditory tasks, participants decided whether the probe and target words were semantically related in a semantic relatedness judgement task. Items were linked by one of two different types of semantic relationships only – taxonomic (i.e. they were in the same semantic category) or thematic (i.e. the items were commonly found or used together). Taxonomic relationships provide hierarchical similarity structures, based primarily on common features shared across same-category items (Hampton, 2006), while thematic relations are largely built around events or scenarios (Estes et al., 2011, Lin and Murphy, 2001), which become conventionalized due to the frequent co-occurrence of particular objects in real-life situations and their linguistic descriptions. On half of the trials, participants were told in advance which relationship would be probed before the presentation of the word pair. For the other half of the trials, participants decided about semantic relatedness based on the two items presented, with no specific instructions in advance. A 2 (Task Knowledge: Known Goal vs. Unknown Goal) × 2 (Semantic Relation: Taxonomic relation vs. Thematic relation) fully-factorial within-subjects design was used to create four conditions, with each experimental condition including 30 related trials. 60 unrelated word pairs were generated without repeating words from the related pairs (i.e., each pair was unique and there was no overlap across conditions). Overall, 120 related and 60 unrelated word pairs were included in this task. The trials were then evenly divided into two sets corresponding to the Known Goal (60 related trials, 30 unrelated) and Unknown Goal (60 related trials, 30 unrelated) conditions.

The assignment of words to conditions was confirmed using an independent sample of 30 participants who provided subjective ratings of thematic relatedness (co-occurrence), taxonomic relatedness (physical similarity), and the difficulty of identifying a connection between the items. Thematically-related word pairs had higher co-occurrence compared to taxonomically-related word pairs, while the taxonomically-related word pairs had higher physical similarity than the thematically-related word pairs. Rated difficulty was the same across both the taxonomic and thematic conditions, and across Known Goal and Unknown Goal trials. For unrelated word pairs, another 12 participants rated Co-occurrence, Physical similarity and Difficulty to confirm the lack of semantic links and equivalence across the Known and Unknown Goal conditions. In addition, linguistic properties (i.e., word frequency, length, and imageability) of the probe and target words were matched across conditions (see Zhang et al., 2021 for details). Word2vec was also used to provide a metric of strength of association for each word pair, since Zhang et al. (2021) included this variable as a parametric regressor to investigate the effects of controlled retrieval demands when linking together more weakly related concepts. Word2vec is a measure of semantic distance that is based on the assumption that words with similar meanings occur in similar contexts (Mikolov, Chen, Corrado, & Dean, 2013), and this metric can capture both taxonomic (physical similarity) and thematic (occurrence in similar contexts) relationships (see Zhang et al., 2021 for details).

The auditory task employed the same words as the published visual task to allow their direct comparison (Zhang et al., 2021). All words in the auditory task (including instruction words) were recorded in a female voice using Praat software (www.fon.hum.uva.nl/praat/). Each audio word was supplied as a 16-bit mono WAV file at a sampling rate of 44100 Hz, and the intensity of each word was processed uniformly to 70dB. The volume of the spoken words output from the headphones was set to a safe level.

Both the visual and auditory scanning sessions also included a non-semantic baseline task. In the visual session, participants were presented with a pair of meaningless letter strings in succession and were asked to decide if they contained the same number of letters (full details in Zhang et al., 2021). In the auditory session, one spoken number was presented as the probe, followed by two spoken numbers in succession. Participants were asked to decide whether the probe number was equal to the sum of the other two numbers. In both the visual and auditory sessions, there were 30 matching trials and 15 mismatching trials in the baseline task.

### 2.4. Procedure

In both the auditory and visual sessions, participants were asked to decide if the words in each pair were semantically related (i.e., either from the same category or thematically related) or unrelated. We manipulated the opportunity for top-down controlled semantic retrieval by changing the instructions before the word pair was presented. On half of the trials, participants were presented with a specific task instruction (‘Category?’ or ‘Thematic?’) before the word pair, so that they knew in advance which type of semantic relationship would be relevant on the trial (Known Goal). On the other half of the trials, participants heard a non-specific instruction (‘Related?’) ahead of the word pair, and they made the decision only from the stimuli alone. The manipulation of task knowledge (i.e., Known Goal vs. Unknown Goal) allowed us to examine whether there is a different response in the brain when participants knew the type of semantic relation between the words in advance or not. In the non-semantic baseline condition, participants in the visual session saw the instruction ’Letter Number’ followed by two meaningless letter strings, while participants in the auditory session heard ‘Equals?’ as the task instruction, followed by a spoken number (for example, ‘9’) as the probe, and two spoken numbers presented in quick succession (for example, ‘1, 8’) as the target to sum (see Figure 1A and 1B for details).

**Figure 1:**
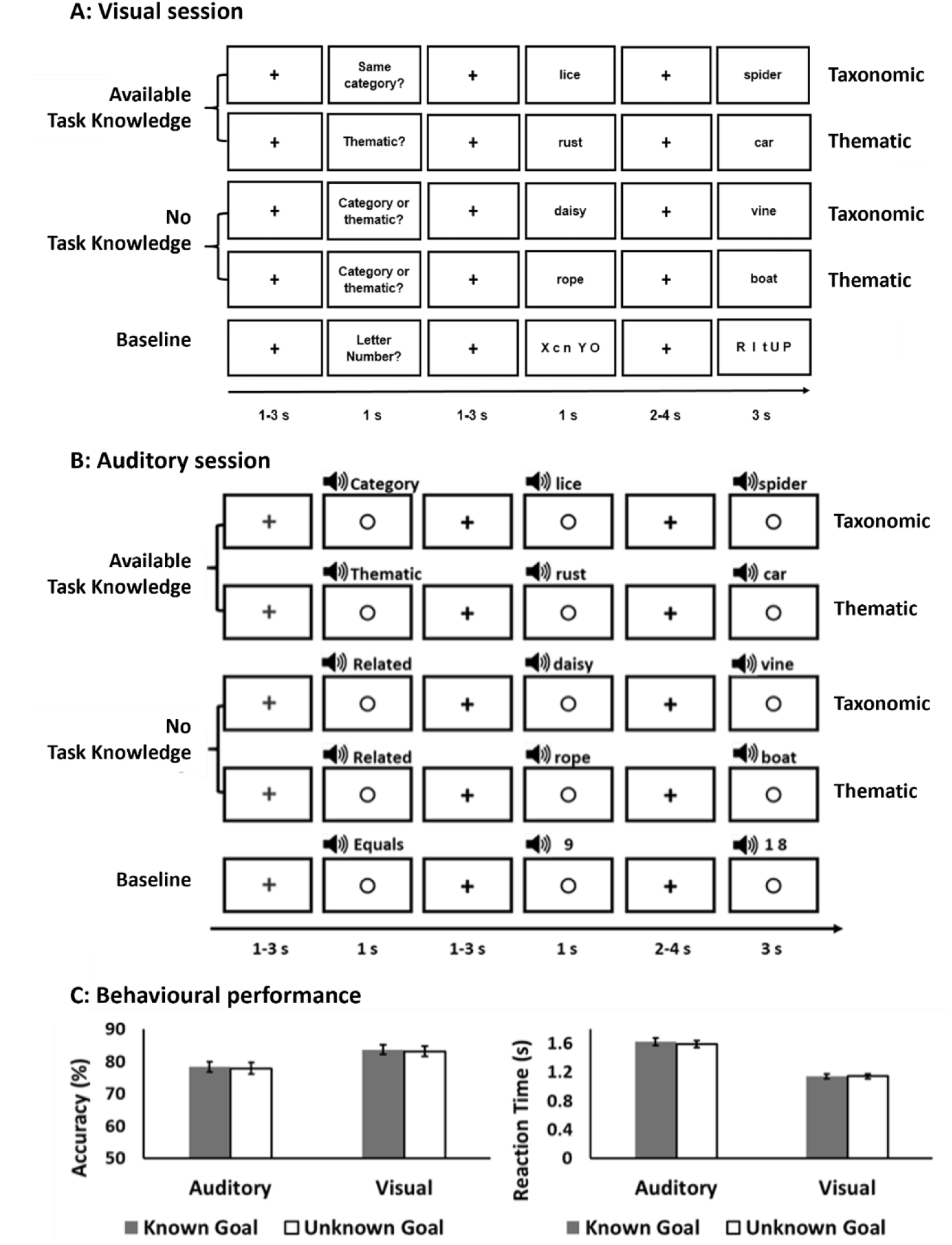
Task design and behavioural data. A: Illustration of visual task, adapted from Zhang, et al. (2021); B: Illustration of auditory task. C: Behavioural results (mean accuracy and reaction time) for auditory and visual modalities. Error bar represents mean standard error.

Figure 1A and 1B indicate the procedure for each condition; this was consistent across written and spoken sessions (Zhang et al., 2021). The stimulus onset timings were held constant across visual and auditory sessions; all audio words (including task instructions, probes and targets) were shorter than one second, and were played synchronously with a visual cue (a circle ‘O’ in the centre of the screen). Each trial started with a red fixation cross presented for a jittered interval of 1–3 s in the centre of the screen. Then the task instruction was presented, followed by a jittered inter-stimulus fixation for 1–3 s. Then the probe item was presented. After a longer jittered fixation interval lasting 2–4 s, the target item was presented during a 3s decision window. Participants pressed one of two buttons on a response box to indicate whether the words were semantically related (or to respond to the non-semantic baseline) as fast and accurately as possible. They used their right index and middle fingers to indicate YES and NO responses. In the auditory condition, if they could not hear the word pair clearly, they pressed a button with their ring finger to indicate this. The ratio of YES to NO responses was held constant across experimental conditions. In both sessions, stimuli were presented in five runs each containing 45 trials: 6 related and 3 unrelated trials in each of the four experimental conditions, and 9 non-semantic baseline trials. Each run lasted 9 minutes, and trials were presented in a random order. The runs were separated by a short break and started with a 9-second alerting slide (i.e., Experiment starts soon).

### 2.5. Neuroimaging data acquisition

Structural and functional data were acquired using a 3T GE HDx Excite Magnetic Resonance Imaging (MRI) scanner utilising an eight-channel phased array head coil at the York Neuroimaging Centre, University of York. Structural MRI acquisition was based on a T1-weighted 3D fast spoiled gradient echo sequence (repetition time (TR) = 7.8 s, echo time (TE) = minimum full, flip angle = 20°, matrix size = 256 × 256, 176 slices, voxel size = 1 mm × 1 mm × 1 mm). Task-based activity was recorded in the same way in the visual and auditory sessions. We used single-shot 2D gradient-echo-planar imaging sequence with TR = 3 s, TE = minimum full, flip angle = 90°, matrix size = 64 × 64, 45 slices, and voxel size = 3 mm × 3 mm × 3 mm. Data for each modality was acquired in a single session. The task was presented across 5 functional runs, each containing 185 volumes.

In an independent sample, a 9-minute resting-state fMRI scan was recorded using single-shot 2D gradient-echo-planar imaging (TR = 3 s, TE = minimum full, flip angle = 90°, matrix size = 64 × 64, 60 slices, voxel size = 3 mm × 3 mm × 3 mm, 180 volumes). The participants were instructed to focus on a fixation cross with their eyes open and to keep as still as possible, without thinking about anything in particular. These data have been used in previous studies (e.g., Evans et al., 2020; Gonzalez Alam et al., 2019; Shao et al., 2022; Vatansever et al., 2017; Wang, X. et al., 2018).

### 2.6. Pre-processing of task-based fMRI data

All functional and structural data were pre-processed using a standard pipeline and analysed via the FMRIB Software Library (FSL version 5.0, www.fmrib.ox.ac.uk/fsl). Individual T1-weighted structural brain images were extracted using FSL’s Brain Extraction Tool (BET). Structural images were linearly registered to the MNI152 template using FMRIB’s Linear Image Registration Tool (FLIRT). The first three volumes (i.e., the presentation of the 9-second task reminder ‘Experiment starts soon’) of each functional scan were removed to minimise the effects of magnetic saturation. The functional data were analysed by using FSL’s FMRI Expert Analysis Tool (FEAT). We applied motion correction using MCFLIRT (Jenkinson, Bannister, Brady, & Smith, 2002), slice-timing correction using Fourier space time-series phase-shifting (interleaved), spatial smoothing using a Gaussian kernel of FWHM 6 mm, and high-pass temporal filtering (sigma = 100 s) to remove temporal signal drift. In addition, motion scrubbing (using the fsl_motion_outliers tool) was applied to exclude volumes that exceeded a framewise displacement threshold of 0.9.

### 2.7. Univariate analysis of task-based fMRI data

The univariate analysis examined the controlled retrieval of semantic information at different stages of the task. The general linear model (GLM) was constructed in an identical fashion for the visual and auditory modalities, following Zhang et al. (2021), to allow a direct comparison of these sessions. We differentiated the response to the first word, when participants accessed semantic representations and prepared for either a specific type of semantic relationship (Specific goal: ‘Category?’ or ‘Thematic?’) or a judgement of either type (Non-specific goal: ‘Related?’), from the second word when participants made a decision about whether the words were linked according to the specific instructions or in the absence of a specific instruction about the nature of the semantic relationship. Consequently, the model included three factors: (1) Word Position (First word vs. Second word), (2) Task Knowledge (Known Goal vs. Unknown Goal), and (3) Semantic Relation (Taxonomic relation vs. Thematic relation). In addition, we included word2vec (Mikolov et al., 2013) as a parametric regressor in the model to characterise the semantic distance between the two words in each trial, in line with the previous study employing written words (Zhang et al., 2021). There were no effects of word2vec at the whole-brain level in the current dataset employing spoken words.

The pre-processed time-series data were modelled using a general linear model, using FMRIB’s Improved Linear Model (FILM) correcting for local autocorrelation (Woolrich, Ripley, Brady, & Smith, 2001). 10 Explanatory Variables (EV) of interest and 5 of no interest were modelled using a double-Gaussian hemodynamic response gamma function (probe and target were modelled separately, therefore there were two EVs for each condition). The 10 EVs of interest were: (1) Probe and (2) Target for Known Goal Taxonomic Relations, (3) Probe and (4) Target for Unknown Goal Taxonomic Relations, (5) Probe and (6) Target for Known Goal Thematic Relations, (7) Probe and (8) Target for Unknown Goal Thematic Relations, (9) Probe and (10) Target for baseline condition (for matching trials, since the semantic data only included related trials). Our EVs of no interest were: (11) Probe and (12) Target for unrelated word pairs, (13) Other inputs-of-no-interest (including the audio instruction word, the period after the response on each trial, and mismatching baseline trials), (14) Fixation (including the inter-stimulus fixations between instructions and first item, as well as between first item and second item when some retrieval or task preparation was likely to be occurring), and (15) Incorrect Responses (including all the incorrect trials across conditions). We also included an EV to model word2vec as a parametric regressor, therefore we had 16 EVs in total. EVs for the first item in each pair commenced at the onset of the audio word or number, with EV duration set as the presentation time (1 s). EVs for the second item in each pair commenced at the onset of the written or spoken word or number, and ended with the participants’ response (i.e. a variable epoch approach was used to remove effects of time on task). The remainder of the second item presentation time was modelled in the inputs-of-no-interest EV (i.e., in the visual task, word 2 was on screen for 3 s, and if a participant responded after 2 s, the post-response period lasting 1 second was removed and placed in the inputs-of-no-interest EV). The parametric word2vec EV had the same onset time and duration as the EVs corresponding to the second word in the semantic trials, but included the demeaned word2vec value as a weight. The fixation period between the trials provided the implicit baseline.

The five sequential runs were combined using fixed-effects analyses for each participant. In the higher-level analysis at the group level, the combined contrasts were analysed using FMRIB’s Local Analysis of Mixed Effects (FLAME1), with automatic outlier de-weighting (Woolrich, 2008). A 50% probabilistic grey-matter mask was applied. Clusters were thresholded using Gaussian random-field theory, with a cluster-forming threshold of z = 3.1 and a familywise-error-corrected significance level of p =.05.

In addition to contrasts examining the main effects of Word Position (Word 1 vs. Word 2), and Task Knowledge (Known Goal vs. Unknown Goal) for both modalities, we included two-way interaction terms of Modality by Task Knowledge, as well as Modality by Word Position. We also included contrasts of all task conditions between visual and auditory modalities (i.e., Known Goal Thematic; Known Goal Taxonomic; Unknown Goal Thematic; Unknown Goal Taxonomic comparing spoken and written words).

### 2.8. Selection of ROIs: semantic control sites and MDN network

Additional ROI analyses characterised the effects of word position and task knowledge in *a priori* semantic control and domain-general executive networks. We examined the SCN using a binarised mask from a recent meta-analysis of semantic control (Jackson, 2021). Domain-general executive regions of the MDN were identified as regions showing difficulty effects across multiple tasks (map taken from Fedorenko et al., 2013). These functionally-defined control networks are partially overlapping, especially in the left hemisphere, although they also have distinct elements – with semantic control extending to more anterior and ventral parts of left inferior frontal gyrus, and left posterior temporal cortex, while MDN is focused on bilateral inferior frontal and intraparietal sulcus. We created four functional network masks from the combination of these network maps: i) MDN regions also implicated in semantic control (i.e., SCN ∩ MDN), ii) SCN-only regions not implicated in domain-general control, iii) MDN-only regions not implicated in semantic control in the left hemisphere, iv) MDN-only regions in the right hemisphere. These maps are available on Neurovault (https://neurovault.org/collections/14750/). In a supplementary analysis, we also present results for each of the five strongest clusters in the SCN meta-analysis (Jackson, 2021).

### 2.9. Spatial correlation between individual-level maps from the univariate analysis and whole-brain cortical gradients

We took non-thresholded maps of each task condition (contrasted against the implicit baseline) at the individual level, and computed the spatial correlation of these maps with the first three cortical gradients derived from intrinsic connectivity (group maps taken from Margulies et al., 2016). This method provides a metric denoting how similar each participant’s pattern of activation and deactivation is to key functional dimensions of cortical organisation. Higher positive spatial correlations with Gradient 1 suggest functional responses that are more heteromodal, while negative correlations mean greater similarity with unimodal regions. Repeated-measures ANOVAs were used to examine whether there were effects of task knowledge, word position or modality on the location of task activation in gradient space.

## 3. Results

### 3.1 Behavioural results

Trials with incorrect responses were excluded from the RT analysis (16.3% in the visual modality; 19.1% reported as not heard clearly and 14.5% incorrect responses on clear trials in the auditory modality). Repeated-measures ANOVAs were performed on both accuracy and RT, examining the effects of Task Knowledge (Known Goal vs. Unknown Goal) and modality (visual vs. auditory)^1^. Figure 1B shows that the auditory task had lower accuracy (left panel, *F*(1, 56) = 6.96, *p* = .011, *η_p_^2^* = .11) and longer reaction times (right panel, *F*(1, 56) = 71.47, *p* < .001, *η_p_^2^* = .56). There were no differences in performance for Known Goal versus Unknown Goal trials (Accuracy: *F*(1, 56) = .24, *p* = .63, *η_p_ ^2^* = .004; RT: *F*(1, 56) = 2.39, *p* = .13, *η_p_ ^2^* = .04) and no interaction (Accuracy: *F*(1, 56) = .01, *p* = .94, *η_p_ ^2^* = .00; RT: *F*(1, 56) = 2.30, *p* = .14, *η_p_^2^* = .04). These behavioural results suggest that while the effect of task knowledge (i.e. top-down control) was equivalent for the two modalities, the auditory task was harder overall because the individual items were more difficult to recognise. Therefore, two types of controlled processing may occur within this task: task knowledge is thought to modulate participants’ capacity to control semantic retrieval in a top-down fashion to suit the task demands (Zhang et al., 2021), while individual spoken words are harder to recognise than written words from the input signal.

### 3.2 Univariate analysis

#### 3.2.1 Activation for auditory and visual semantic decisions

First, we examined the brain regions engaged in the semantic decision-making phase in a whole-brain analysis, in the visual and auditory modalities respectively. We conducted a conjunction analysis to investigate the regions recruited in common across written and spoken words (Figure 2). Semantic decisions to written words (i.e., significant clusters for the contrast of Word 2 > Word 1 for visually presented words) included bilateral inferior frontal gyrus (IFG), prefrontal cortex, occipital pole, left inferior temporal gyrus (ITG) and precentral cortex. Semantic decisions to spoken words (i.e., significant clusters for the contrast of Word 2 > Word 1 in the auditory domain) included left IFG, lateral occipital cortex and precentral cortex. A conjunction of these maps for written and spoken words showed common activation across modalities in left IFG and dorsolateral prefrontal cortex, pre-supplementary motor area and posterior cingulate cortex.

**Figure 2:**
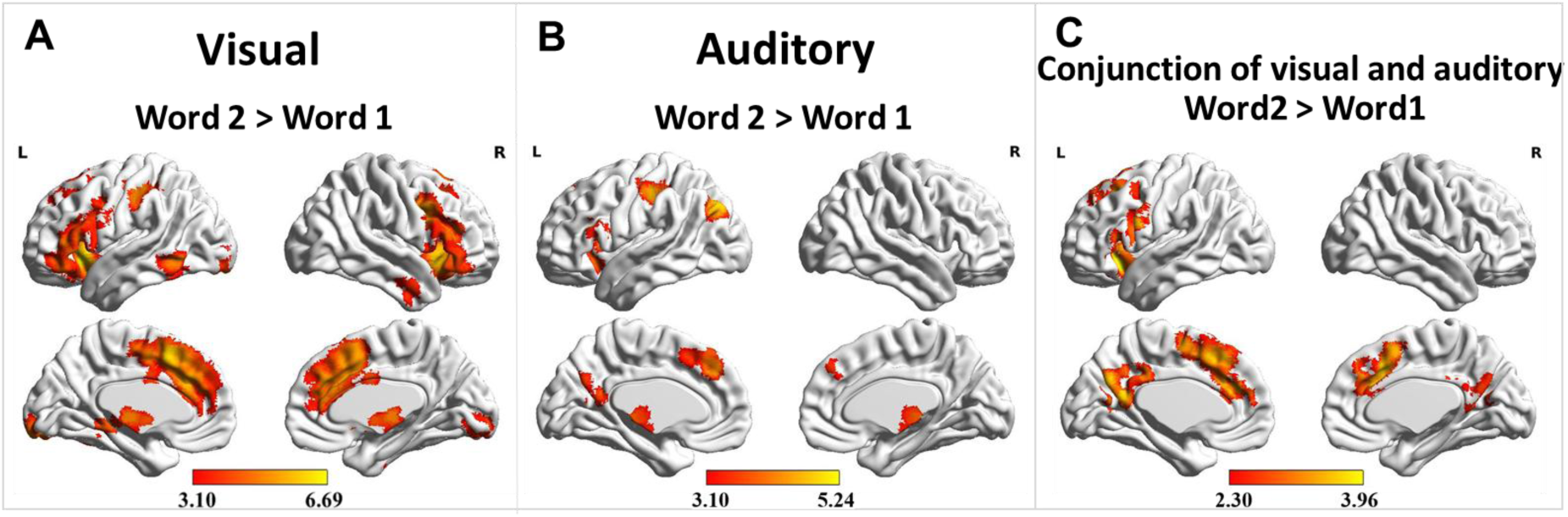
Effect of word position (i.e., the effect of decision-making) in the visual (Figure 2A) and auditory modality (Figure 2B), and regions activated during semantic decisions for both modalities (Figure 2C). Maps in Figure 2A and 2B were cluster-corrected with a voxel inclusion threshold of z > 3.1 and family-wise error rate using random field theory set at p <lt; .05. The conjunction map in Figure 2C is identified using FSL’s ‘easythresh_conj’ tool and thresholded at z = 2.3.

The global contrast between visual and auditory tasks, regardless of word order, is shown in Supplementary Figure S1. Spoken words elicited more activation in and around bilateral auditory cortex and supplementary motor area. Written words elicited more activation in posterior ITG and fusiform cortex close to the visual word form area, posterior intraparietal sulcus, frontal eye fields and frontal pole.

#### 3.2.2 The interaction between word position and modality

Next, we considered whole-brain differences between auditory and visual modalities in the effect of (i) word position (contrasts of words 1 and 2) and (ii) task knowledge. These analyses can establish if there are differences in the neural basis of semantic decision-making and top-down semantic control when comparing spoken and written words.

The interaction of modality with word position revealed three clusters, in left ventral posterior temporal cortex, left occipital pole, and right inferior frontal gyrus (Figure 3A). To examine the role of each cluster, we determined their resting-state functional connectivity and functionally decoded these unthresholded maps using Neurosynth (Yarkoni et al., 2011). The top 10 relevant terms corresponding to the three clusters were chosen and rendered as word clouds (shown in Figure 3B). Ventral posterior temporal cortex (cluster 1 in Figure 3A) was activated by both written and spoken words, but the effect of word position was stronger for the visual task. Text and speech showed opposite patterns, with written words showing more activation during decision-making (word 2 > word 1), and spoken words showing more activation for the first word in the pair (when word recognition was more challenging). The connectivity pattern of this cluster was related to terms such as ‘working memory’, ‘word’ and ‘semantic’.

**Figure 3.**
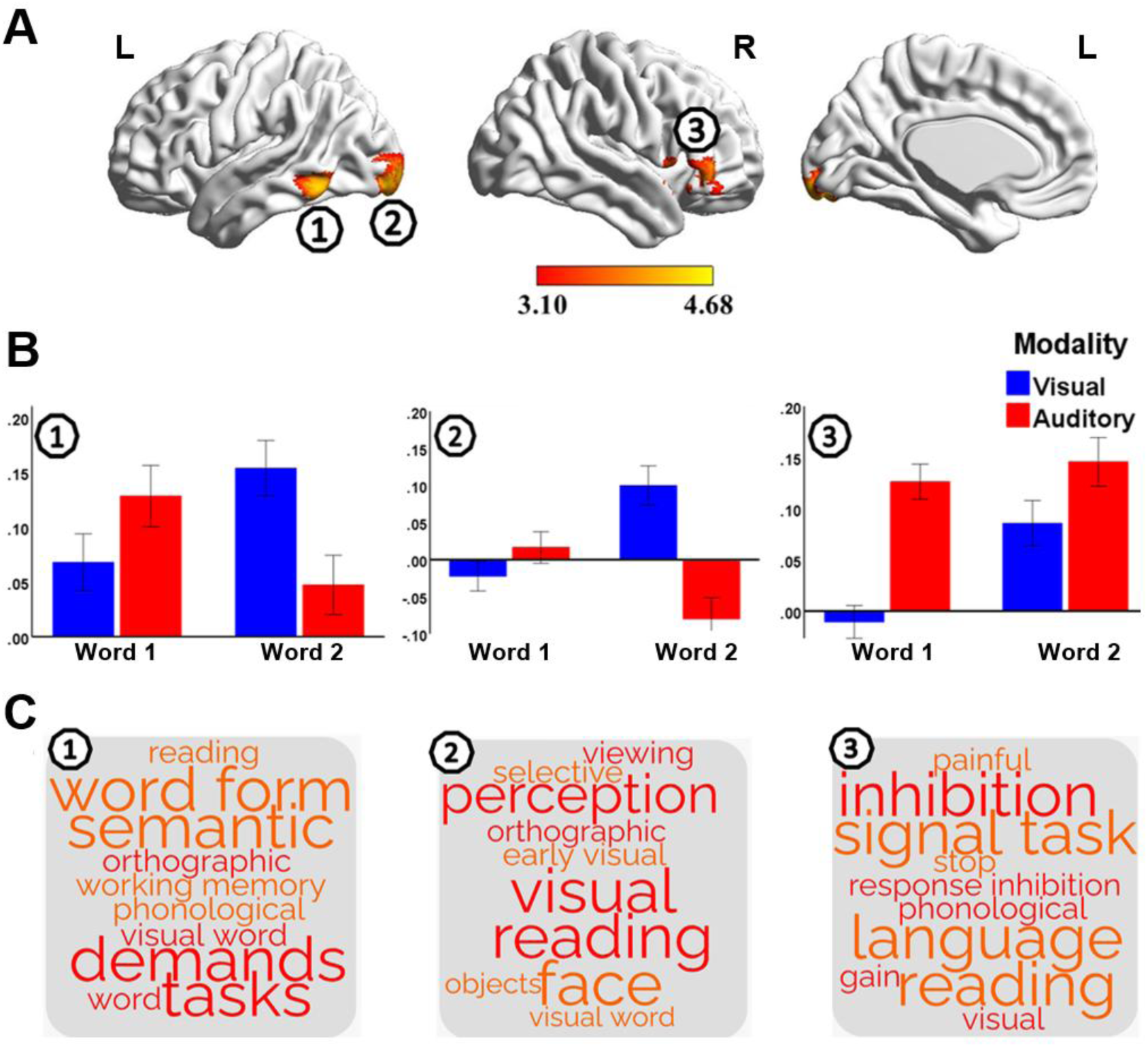
Differences between modalities in the effect of word position. Three clusters (Figure 3A) were identified in this whole-brain analysis of the interaction between modality and word position; maps are cluster-corrected at a threshold of *z* > 3.1 (FWE corrected *p* <lt; .05). Decoding was performed on the unthresholded intrinsic connectivity maps seeded from the three clusters, and presented in a word cloud for each cluster in Figure 3B. The signal change within each cluster is shown in Figure 3C.

Occipital pole (cluster 2 in Figure 3) showed relatively little response to the first word of the pair in either modality but activated to visual decision-making, while deactivating in response to auditory decision-making. The connectivity of this cluster was associated with visual processing and reading.

Finally, the cluster in right IFG (cluster 3 in Figure 3) showed a stronger response to the auditory task on the first word, when the item was hardest to recognise. It then responded to the second word across both modalities. The connectivity pattern of this cluster was associated with both executive and language processes. These results might therefore reflect stronger engagement of control mechanisms in both auditory speech recognition, and in semantic decision-making.

#### 3.2.3 The interaction between task knowledge and modality

Next, we examined the interaction of modality with task knowledge, revealing differences across visual and auditory inputs in the effect of prior knowledge about the nature of the semantic relationship to be retrieved. Effects of task knowledge for visual inputs were identified in left IFG by Zhang et al. (2021; yellow in Figure 4A). However, we did not observe similar effects for the auditory modality, and there were no main effects of task knowledge in the whole-brain analysis. There was a modality by task knowledge interaction in a cluster bordering the left IFG and insula (the red cluster in Figure 4A; signal change within this cluster is shown in Figure 4C), which reflected a stronger effect of task knowledge in the visual domain. We used Neurosynth (Yarkoni et al., 2011) to decode the unthresholded connectivity maps of the visual task knowledge effect (yellow in Figure 4A) and the modality by task knowledge interaction (red in Figure 4A). The connectivity maps are available on Neurovault (https://neurovault.org/collections/14750/). The top 10 terms obtained from cognitive decoding of the connectivity patterns of these clusters revealed terms such as ‘word’ and ‘semantic’ for Zhang et al.’s task knowledge main effect for written words (yellow in Figure 4A) and ‘motor’ and ‘movement’ for the interaction of modality and task knowledge (red in Figure 4A). In the interaction cluster, written words showed a stronger response when task knowledge was available, while spoken words showed a stronger response without task knowledge. This might reflect differences in the source of difficulty across these two modalities: for the auditory modality there is more uncertainty about the identity of individual words, especially in the absence of task knowledge, while for written words, the most difficult aspect of the task involves establishing a semantic link between the words. Both of these aspects of controlled semantic cognition could involve this region implicated in motor processing.

**Figure 4:**
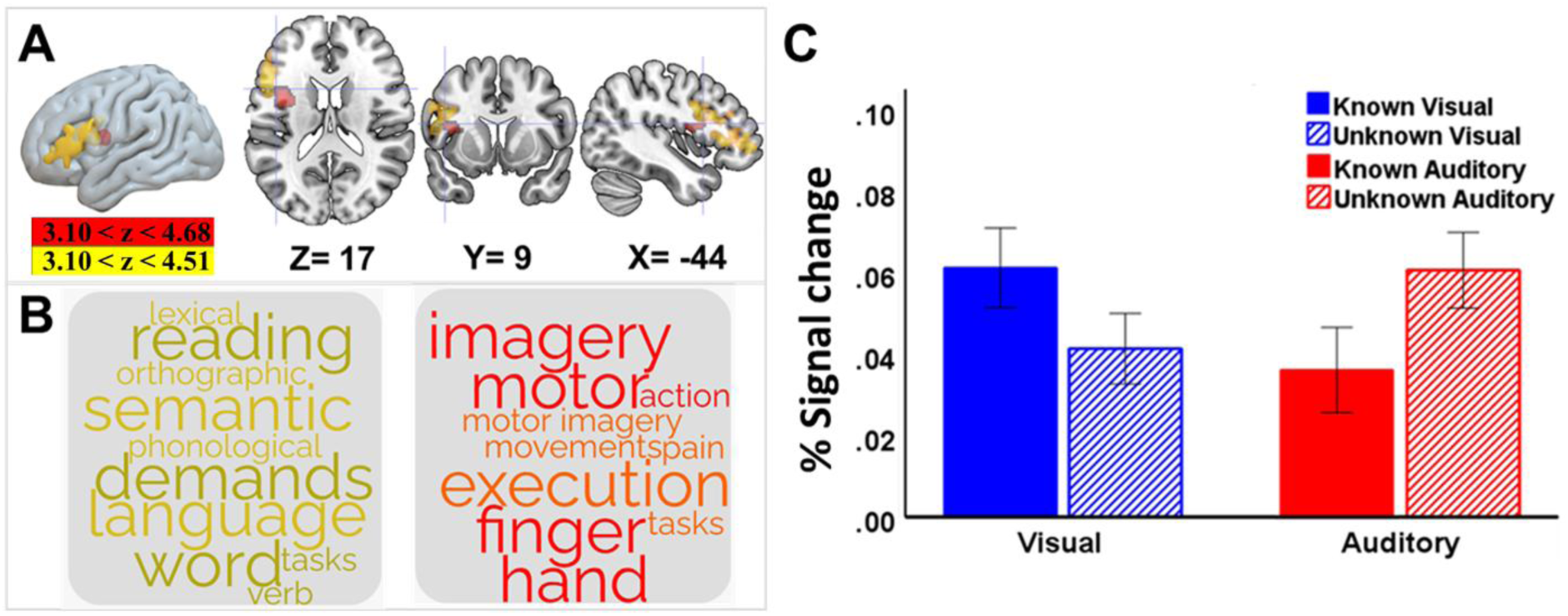
Differences between modalities in the effect of task knowledge. One cluster bordering left posterior insula and IFG (cluster in red colour in Figure 4A) was identified in a whole-brain interaction between modality and task knowledge. This interaction effect was adjacent to the effect of task knowledge in the visual modality (Known Goal > Unknown Goal; yellow in Figure 4A). All maps are cluster-corrected at a threshold of *z* > 3.1 (FWE corrected *p* <lt; .05). Decoding was performed on the unthresholded maps reflecting the effect of task knowledge in the visual modality (Figure 4A, yellow) and the interaction between modality and task knowledge (Figure 4A red). Figure 4C shows the signal change within the interaction cluster (red cluster in Figure 4A) in written and spoken tasks respectively. Error bars represent mean standard error.

The whole-brain interactions between modality and word position/task knowledge suggest there are different kinds of task demands operating in our paradigm. Spoken words were more difficult to recognise, and this effect is expected to be greater in the absence of any context (e.g., for the first word and without task knowledge). This might explain stronger activation for the auditory task in clusters linked to cognitive control, including in left ventral posterior temporal cortex and right inferior frontal gyrus (for word 1) and in left posterior insula/inferior frontal gyrus (for the unknown auditory condition). In contrast, we see stronger activation of control sites for visually-presented words during semantic decision-making (for word 2) and when task knowledge is available in advance (known condition); conditions in which participants can particularly benefit from top-down control to establish a semantic link.

### 3.3 Situating the individual-level maps from univariate analysis in the whole-brain principal gradient

To investigate the extent to which *whole-brain* responses to written and spoken words reflect a different balance of processing between abstract/heteromodal and sensory-motor regions, and how these differences depend on the availability of task knowledge, we computed spatial correlations between the unthresholded individual-level contrast maps of each condition (compared with the implicit baseline) and the whole-brain principal gradient (from Margulies et al., 2016), which captures the distinction in connectivity between unimodal and heteromodal regions. The principal gradient is correlated with distance from primary sensory-motor landmarks and describes the sequence of resting-state networks seen along the cortical surface, from sensory-motor, through attention and frontoparietal networks to default mode network in multiple areas of cortex. Positive correlations reveal functional responses that are relatively heteromodal, while negative correlations suggest a stronger response towards unimodal cortex. Correlations between activation maps and the principal gradient were negative overall, reflecting sensory-motor responses elicited by the task relative to baseline; these low-level responses are controlled in the comparisons between conditions below.

An ANOVA of correlations with the principal gradient revealed significant main effects of *Modality* (*F*(1,56) = 11.09, *p* = .002, *η_p_^2^* = .17), *Word Position* (*F*(1, 56) = 6.49, *p* = .014, *η_p_^2^* = .10) and *Task Knowledge* (*F*(1, 56) = 5.77, *p* = .020, *η_p_^2^* = .09), reflecting a more heteromodal response for written than spoken words, for Word 2 versus Word 1, and for Known compared with Unknown Goal trials. The two-way interaction between *Task Knowledge* and *Modality* (*F*(1, 56) =5.70, *p* = .020, *η_p_^2^* = .09) and the three-way interaction (*F*(1, 56) = 7.12, *p* = .010, *η_p_^2^* = .11) were also significant. Follow-up ANOVAs showed effects of *Word Position* for the visual but not the auditory modality. Written words showed a higher correlation with Gradient 1 in the decision-making phase at Word 2 compared with Word 1 (*F*(1, 30) = 5.46, *p* = .026, *η_p_^2^* = .15), and this effect of *Word Position* interacted with *Task Knowledge* (*F*(1, 30) = 6.78, *p* = .014, *η_p_^2^* = .18), reflecting a stronger effect of word position on Known Goal trials. These effects of *Word Position* were not seen in the auditory modality, although there was a main effect of *Task Knowledge* (*F*(1, 26) = 8.21, *p* = .008, *η_p_^2^* = .24) with *Unknown* Goal trials eliciting a stronger response for spoken words towards sensorimotor regions. In this way, analysis of the activation pattern along the principal gradient found that written words elicited a pattern of activation that was less sensory-motor and more heteromodal, especially during decision-making and when task goals were known: these circumstances strengthened the contribution of heteromodal processes to the task state as a whole. In contrast, the balance of processing for spoken words was tilted towards unimodal processes potentially reflecting the relative difficulty of input processing in the auditory task and/or the engagement of control mechanisms focussed on sensory-motor processes.

### 3.4 ROI analysis to characterise effects of task knowledge and word position in SCN and MDN

The analysis above demonstrates that the balance of unimodal to heteromodal processes is influenced by task knowledge in a different way across modalities. The next analysis considers activation within control systems: although both SCN and MDN are typically thought to be ‘heteromodal’, recent research has shown that these networks occupy different positions on the principal gradient: SCN is closer to the heteromodal end of this gradient than MDN (Chiou et al., 2023; Wang et al., 2020), consistent with the abstract and heteromodal nature of semantic processing, and in line with our observation that SCN responds to task knowledge across modalities, while MDN responds during difficult auditory perception.

Given our results suggest different control demands operate in our task – with input processing being more challenging for spoken words (on word 1, in the absence of task knowledge), while top-down control can be applied more readily when task knowledge is available (on word 2) – we tested for these different effects of control within four networks of interest: i) the semantic control network, which is largely left lateralised (SCN, overlap with MDN was excluded), ii) the overlap of SCN and MDN, which is again largely left lateralised (SCN∩MDN), and iii) the regions of the MDN which do not overlap the SCN in the left hemisphere (MDN left) and iv) in the right hemisphere (MDN right).

All sites showed main effects of word position with higher activation for the second word (SCN: *F*(1,56) = 90.94, *p* < .001, *η_p_^2^* = 1.00; SCN∩MDN: *F*(1,56) = 143.62, *p* < .001, *η_p_^2^* = .72; MDN left: *F*(1,56) = 17.44, *p* < .001, *η_p_^2^* = .24; MDN right: *F*(1,56) = 6.70, *p* = .012, *η_p_^2^* = .11) and no interaction between modality and *Word Position*. However, the networks showed different effects of task knowledge. SCN and SCN∩MDN showed main effects of task knowledge (more activation with a Known Goal (SCN: *F*(1,56) = 13.30, *p* < .001, *η_p_^2^* = .95; SCN∩MDN: *F*(1,56) = 10.17, *p* = .002, *η_p_^2^* = .15) and no interaction between modality and task knowledge. MDN (SCN∩MDN and MDN in right hemisphere) showed modality effects with higher activation for the harder auditory task (SCN∩MDN: *F*(1,56) = 8.83, *p* = .004, *η_p_^2^* = .14; MDN right: *F*(1,56) = 4.45, *p* = .039, *η_p_^2^* = .07; this pattern was not found for left MDN). These results suggest that MDN is more important for controlled sensory-motor processing, including the identification of word meaning from spoken inputs in a noisy environment, while SCN is more relevant to top-down semantic control from goals or expectations.

Supplementary Figure S2 shows these effects for the strongest five clusters of the SCN separately (including voxels that also fall within MDN). All sites showed effects of word position with higher activation for Word 2. Most sites (except left pMTG) showed effects of task knowledge with higher activation for Known Goal trials. Effects of modality were inconsistent, with key left-hemisphere nodes in left IFG and pMTG showing a heteromodal response that was equivalent across visual and auditory inputs, and anterior and posterior right IFG and dorsomedial prefrontal cortex (dmPFC), also implicated in domain general control (see Figure 5a), showing a stronger response to spoken words.

**Figure 5.**
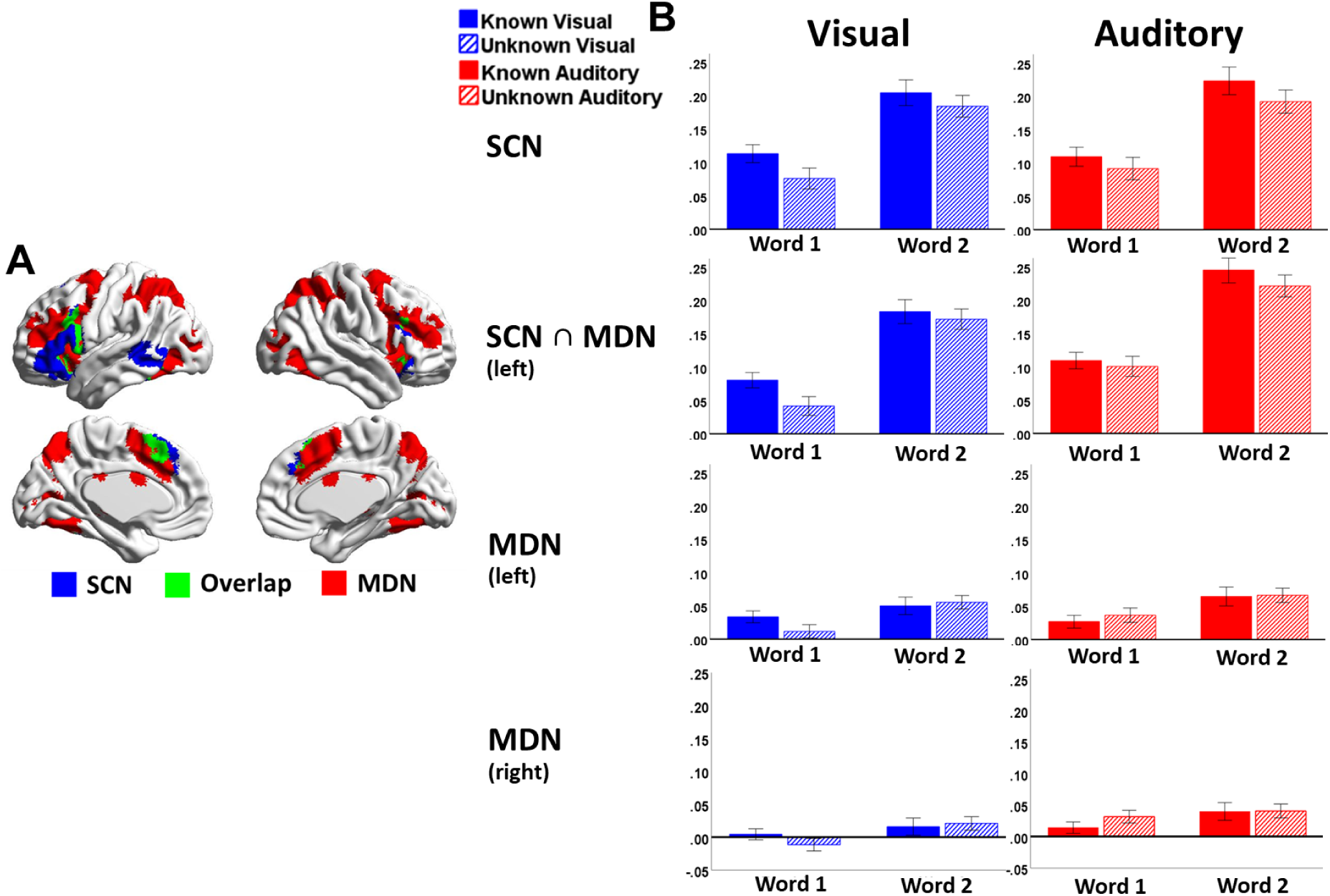
ROI analysis examining the engagement of SCN and MDN. Four ROIs are illustrated in Figure 5A: SCN network (excluded the overlap with MDN, shown in blue), overlap of SCN and MDN (in left hemisphere, shown in green), MDN network (excluding overlap with SCN, shown in red) in left and right hemispheres.

**Figure 6.**
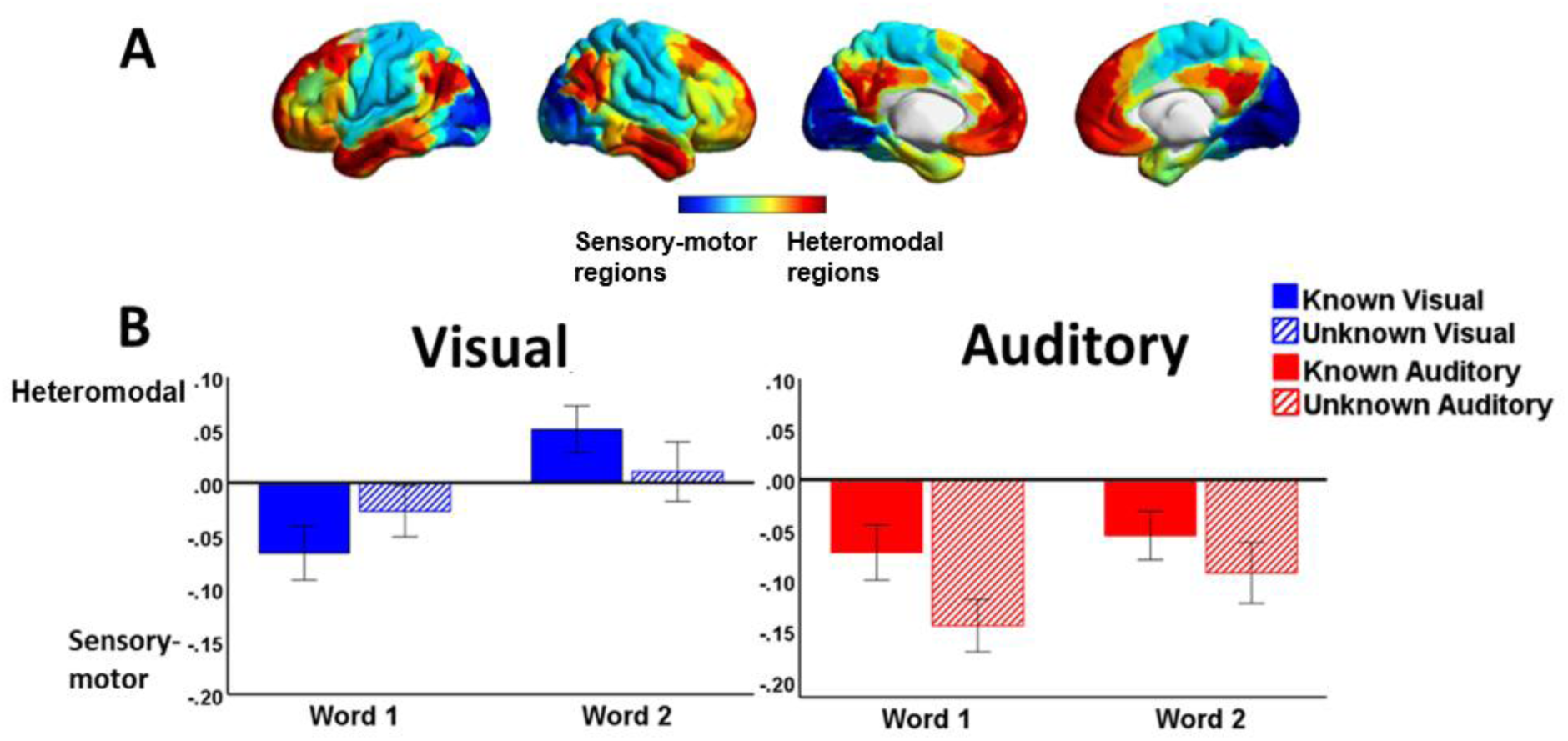
Spatial correlations between the unthresholded individual-level contrast maps and the principal gradient derived from resting-state functional connectivity (Margulies et al., 2016, see Figure 6A). The correlations for the known and unknown conditions at each word position are presented in Figure 6B.

## 4. Discussion

The current study investigated the effect of modality on semantic control by manipulating task knowledge in spoken versus written semantic judgements. The decision-making phase of the task, across both visual and auditory inputs, elicited activation in heteromodal brain areas implicated in cognitive control and semantic processing.

Nevertheless, we identified different kinds of task demands – linked to top-down effects of task knowledge and the bottom-up difficulty of identifying spoken words – which dissociated across control networks relevant to semantic cognition (SCN and MDN).

Clusters in right frontal and left posterior temporal cortex, associated with task demands and executive control, showed a stronger response to the first word of the pair for spoken words, when the inputs were particularly hard to recognise. In contrast, left IFG (commonly implicated in semantic control), showed more activation for visual inputs in the presence of task knowledge, suggesting that this site supports the top-down control of retrieval. These different aspects of task difficulty were also captured by the principal gradient of connectivity: written words elicited more activation towards the heteromodal end of the principal gradient, particularly during semantic decision making when the second word of the pair was presented, and when the goal for semantic retrieval was known in advance.

Under these conditions, semantic control processes towards the DMN end of the principal gradient are expected to play a key role in the neurocognitive state supporting task performance. In contrast, spoken words were associated with activation closer to the unimodal end of the gradient, suggesting that sensory-motor processes make a more important contribution to this task. Finally, we established a dissociation between functional networks implicated in cognitive control: SCN showed a stronger response to Known Goal trials across modalities, consistent with its purported role in constraining semantic retrieval to suit the circumstances (Davey et al., 2016; Jackson 2021; Lambon Ralph et al., 2017; Noonan et al., 2010; Noonan et al., 2013); in contrast, MDN showed more activation to spoken inputs which were harder to recognise, in line with its role in maintaining attention to external goal-relevant stimuli in an effortful fashion in demanding tasks (Brownsett et al., 2014). These findings, taken together, provide comprehensive evidence for dissociable neurocognitive effects underpinning controlled semantic processing.

Our findings are consistent with previous evidence showing that left IFG is involved in the controlled retrieval of semantic knowledge – responding to both bottom-up manipulations of control demands, for example, weak versus strong associations (Badre et al., 2005; Krieger-Redwood et al., 2015; Noonan et al., 2013; Whitney et al., 2011), and top-down instructions to focus on specific features (Chiou et al., 2018; Davey et al., 2016). Zhang et al. (2021) found that left IFG had higher activation in the visual task when the goal for semantic retrieval was known in advance, reflecting top-down controlled processing. The most difficult aspect of this visual task was establishing a semantic link between the words.

In contrast, auditory input is inherently ambiguous; in the current study, an adjacent region of posterior left IFG and insula also showed top-down effects of task knowledge for written words, yet this site was more engaged by spoken words when task knowledge was *not* available, perhaps because control processes supported by this region were recruited to reduce uncertainty about the input (at word 1). Our findings are broadly consistent with the view that there are functional subdivisions within LIFG, with anterior regions supporting semantic control selectively, and more posterior sites contributing to other aspects of controlled language processing (Badre et al., 2005; Badre and D’Esposito, 2009, Demb et al., 1995, Gabrieli et al., 1998). The results also fit with previous evidence revealing that the posterior insula is connected to visual, auditory and sensorimotor cortices, allowing this site to support diverse functions within and beyond language (Zhang et al., 2019; Bamiou, Musiek & Luxon, 2003).

SCN showed stronger activation when it was possible to apply a task goal in a top-down fashion across both visual and auditory inputs, while MDN showed a stronger response in the auditory task which was also harder in terms of behavioural performance. This provides further support for the conclusions of recent studies showing functional dissociations between these networks (Chiou et al., 2023; Gao et al., 2021; Hodgson, Lambon Ralph, & Jackson, 2024): for example, Gao et al. found that SCN responded to controlled semantic retrieval demands, while MDN showed a stronger response to non-semantic language demands (verbal working memory load). SCN is sandwiched between DMN and MDN topographically, and SCN is closer to the heteromodal end of the principal gradient compared to MDN (Chiou et al., 2023; Wang et al., 2020). This topographical organisation appears to relate to the functions of SCN and MDN. SCN supports flexible semantic retrieval, which is inherently more abstract and heteromodal than sensory or motor control; here we show the recruitment of relatively heteromodal semantic control processes supports the ability to apply goal information to conceptual retrieval in a top-down fashion. MDN, which is closer to sensorimotor and attention networks on the principal gradient, showed stronger activation relating to perceptual difficulty. This might suggest MDN is more important for the control of external perception and action than SCN (cf. Wang et al., 2024). Although MDN is largely insensitive to the modality of task presentation, it is adjacent to regions that show visual and auditory preferences (Assem et al., 2022; Wang et al., 2024). Therefore, the principal gradient might organise a hierarchy of regions that support control, extending from areas sensitive to input modality proximal to MDN, through to highly heteromodal responses within SCN near the DMN apex (Chiou et al., 2023).

While these aspects of control were separable both psychologically (in terms of how they were influenced by task manipulations) and anatomically (in terms of differences in activation between SCN and MDN), several clusters associated with cognitive control showed both effects. First, a cluster in temporal-occipital cortex, bordering visual word form area and MDN, supported auditory perception when demands were higher (to the first word in the pair) as well as decision-making to written words, when controlled semantic retrieval is expected to be more engaged. Secondly, left posterior IFG/insula, implicated in motor processing in the cognitive decoding analysis, showed a stronger response to auditory inputs in the absence of task knowledge (when demands on auditory recognition are higher) and to visual inputs in the presence of task knowledge (when top-down semantic control is possible). There are at least two potential explanations for these complex effects: first, both sites are close to both SCN and MDN and might support both aspects of control (or potentially separable effects at the individual level are being merged in our group analysis).

Alternatively, orthographic processes in temporal-occipital cortex and verbal motor processes in posterior IFG/insula might be implicated in both word recognition and semantic decision-making.

Although we utilised the same task paradigm and items for spoken and written words, we examined the effect of modality across two groups of participants; this has the advantage that we can be confident that there is no influence of episodic memory within the modality contrast, but differences in functional organisation between participants might mask fine-grained functional variation attributable to modality. Moreover, performance was not matched across the auditory and visual tasks: participants indicated that they were not always able to hear the spoken words and they look longer to respond to them. While we did not directly manipulate perceptual difficulty for spoken and visual words, the background noise in the MRI scanner would have influenced auditory perception more than visual perception. A direct contrast of the effect of perceptual difficulty across modalities within the same sample would help to confirm whether the stronger activation for auditory inputs in MDN reflected this aspect of difficulty.

Despite these unresolved issues, our study shows that different aspects of semantic task demands dissociate across sites, networks and within the principal gradient. We found a dissociation between task demands associated with the top-down application of task knowledge and with the identification of word meaning from spoken inputs in a noisy environment across SCN and MDN. These results suggest distinct functional roles for cognitive control networks that reflect their location within an axis of functional organisation that extends from unimodal to heteromodal cortex.

## Footnotes

1 The effect of semantic relation (taxonomic vs. thematic) was not examined. This variable was included only to provide a means of manipulating task knowledge prior to retrieval.

