## Supplementary Figures for "The neural processes underpinning flexible semantic retrieval in visual and auditory modalities"

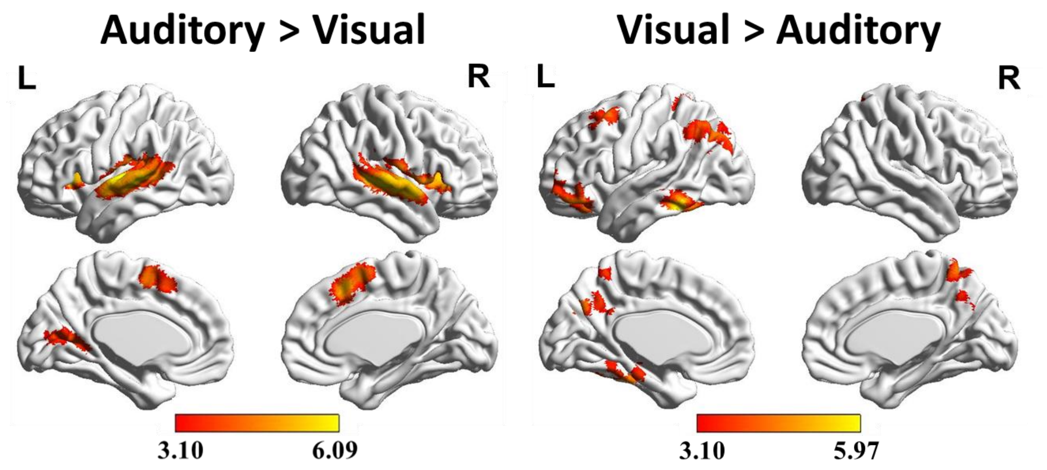


Supplementary Figure S1. The contrast between visual and auditory tasks (i.e., all four task conditions in auditory modality versus all four task conditions in visual modality). Maps were cluster-corrected at a threshold of z > 3.1 (p < .05).

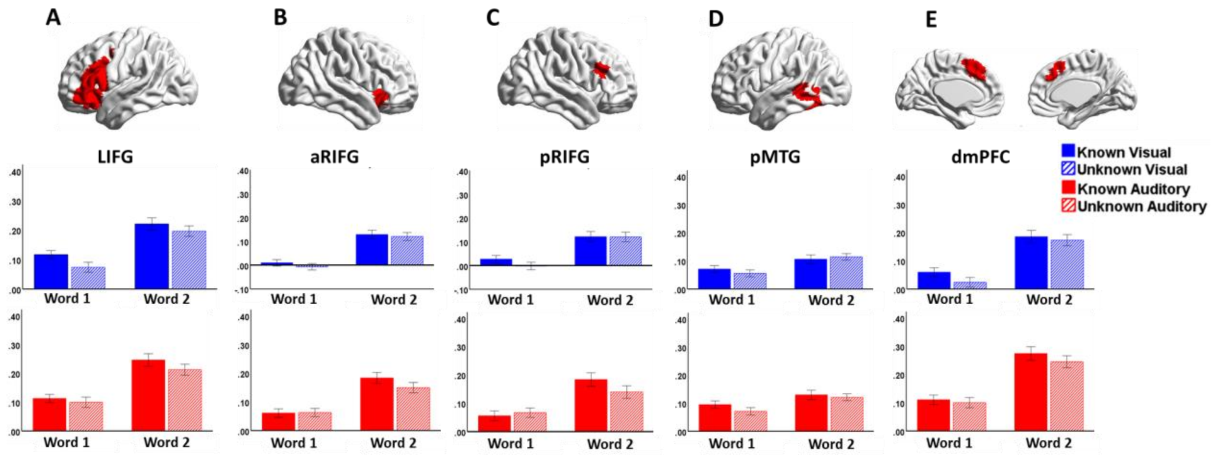


Supplementary Figure S2. Mean signal change in each cluster of the semantic control network (taken from Jackson 2021) for Known and Unknown trials relative to implicit baseline, at the first and second words, in visual and auditory tasks.

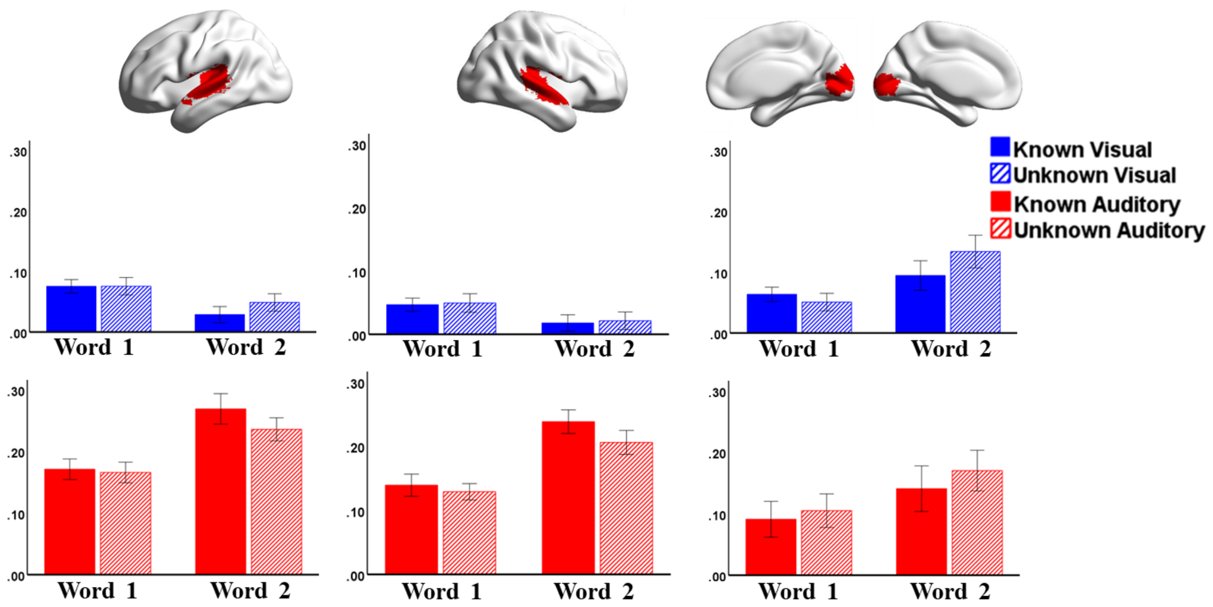


Supplementary Figure S3. Mean signal change in primary visual (one cluster in both left and right hemisphere) and auditory cortex (one cluster in each hemisphere respectively) for Known and Unknown trials relative to implicit baseline, at the first and second words, in visual and auditory tasks. Masks of primary visual and auditory cortex are derived by using the terms ‘primary visual’ and ‘primary auditory’ in NeuroSynth (http://neurosynth.org; Yarkoni et al., 2011).

Supplementary Table S1. Results of repeated-measured ANOVAs for mean signal change in each cluster of the semantic control network (taken from Jackson 2021) for Known and Unknown trials relative to implicit baseline, at the first and second words, in visual and auditory tasks.

| | | LIFG | | | aRIFG | | | pRIFG | | | pMTG | | | dmPFC | | |
|---|---|---|---|---|---|---|---|---|---|---|---|---|---|---|---|---|
| | | F | p | $\eta^2$ | F | p | $\eta^2$ | F | p | $\eta^2$ | F | p | $\eta^2$ | F | p | $\eta^2$ |
| **Main effect** | Modality | F(1,56) = .709 | .403 | .013 | F(1,56) = 10.125 | **.002** | .153 | F(1,56) = 4.749 | **.034** | .078 | F(1,56) =1.413 | .240 | .025 | F(1,56) = 10.666 | **.002** | .160 |
| | Task knowledge | F(1,56) = 13.710 | **< .001** | .197 | F(1,56) = 4.069 | **.048** | .068 | F(1,56) = 5.600 | **.021** | .091 | F(1,56) = 2.322 | .133 | .040 | F(1,56) = 8.144 | **.006** | .127 |
| | Word position | F(1,56) = 101.071 | **< .001** | .64 | F(1,56) = 91.26 | **< .001** | .620 | F(1,56) = 58.43 | **< .001** | .511 | F(1,56) = 31.102 | **< .001** | .357 | F(1,56) = 132.541 | **< .001** | .703 |
| **Two-way interaction** | Modality × Task Knowledge | F(1,56) = .470 | .496 | .008 | F(1,56) = .047 | .828 | .001 | F(1,56) = .005 | .946 | .000 | F(1,56) = 1.020 | .317 | .018 | F(1,56) = .089 | .766 | .002 |
| | Modality × Word position | F(1,56) = .193 | .662 | .003 | F(1,56) = .650 | .424 | .011 | F(1,56) = .082 | .776 | .001 | F(1,56) = .065 | .800 | .001 | F(1,56) = .471 | .495 | .008 |
| | Task knowledge × Word position | F(1,56) = .003 | .957 | .000 | F(1,56) = .628 | .432 | .011 | F(1,56) = .553 | .460 | .010 | F(1,56) = 2.340 | .132 | .040 | F(1,56) = .034 | .853 | .001 |
| **Three-way interaction** | Modality × Task Knowledge × Word position | F(1,56) = 1.245 | .269 | .022 | F(1,56) = 1.638 | .206 | .028 | F(1,56) = 4.892 | **.031** | .080 | F(1,56) = .059 | .809 | .001 | F(1,56) = 1.811 | .184 | .031 |

Supplementary Table S2. Results of repeated-measured ANOVAs for mean signal change in primary visual (one cluster in both left and right hemisphere) and auditory cortex (one cluster in each hemisphere respectively) for Known and Unknown trials relative to implicit baseline, at the first and second words, in visual and auditory tasks.

| | | **Left primary auditory cortex** | | | **Right primary auditory cortex** | | | **Primary visual cortex** | | |
|---|---|---|---|---|---|---|---|---|---|---|
| | | F | p | $\eta^2$ | F | p | $\eta^2$ | F | p | $\eta^2$ |
| **Main effect** | Modality | F(1,56) = 38.193 | **< .001** | .405 | F(1,56) = 48.527 | **< .001** | .464 | F(1,56) = .627 | .432 | .011 |
| | Task knowledge | F(1,56) = .184 | .670 | .003 | F(1,56) = 1.260 | .266 | .022 | F(1,56) = 3.530 | .065 | .059 |
| | Word position | F(1,56) = 1.501 | .226 | .026 | F(1,56) = 3.717 | .059 | .062 | F(1,56) = 8.238 | **.006** | .128 |
| **Two-way interaction** | Modality × Task Knowledge | F(1,56) = 6.069 | **.017** | .098 | F(1,56) = 3.816 | .056 | .064 | F(1,56) = .130 | .072 | .002 |
| | Modality × Word position | F(1,56) = 21.348 | **< .001** | .276 | F(1,56) = 22.207 | **< .001** | .284 | F(1,56) = .215 | .644 | .004 |
| | Task knowledge × Word position | F(1,56) = .007 | .933 | .000 | F(1,56) = .476 | .493 | .008 | F(1,56) = 4.623 | **.036** | .076 |
| **Three-way interaction** | Modality × Task Knowledge × Word position | F(1,56) = .872 | .354 | .015 | F(1,56) = .587 | .447 | .010 | F(1,56) = 2.014 | .161 | .035 |
